# Fourier spectral density of the coronavirus genome

**DOI:** 10.1101/2020.06.30.180034

**Authors:** H.S. Tan

## Abstract

We present an analysis of the coronavirus RNA genome via a study of its Fourier spectral density based on a binary representation of the nucleotide sequence. We find that at low frequencies, the power spectrum presents a small and distinct departure from the behavior expected from an uncorrelated sequence. We provide a couple of simple models to characterize such deviations. Away from a small low-frequency domain, the spectrum presents largely stochastic fluctuations about fixed values which vary inversely with the genome size generally. It exhibits no other peaks apart from those associated with triplet codon usage. We uncover an interesting, new scaling law for the coronavirus genome: the complexity of the genome scales linearly with the power-law exponent that characterizes the enveloping curve of the low-frequency domain of the spectral density.

## 1 Introduction

Motivated by our search for deeper organizational principles governing genetic information ^1^, the study of a DNA/RNA genome via its Fourier spectral density has given us several interesting insights into the code of life. An example of a seminal paper in this subject is that of Voss in [2] where the author found that the spectral density of the genome of many different species follows a power law of the form 1*/k*^*β*^ in the low-frequency domain, with the exponent *β* potentially related to the organism’s evolutionary category. In [2], *β* was found to be close to 1, a phenomenon shared by a wide variety of physical systems especially those that carry long-range correlations or characterized by a myriad of length scales. It was also found that the power spectra may contain defining peaks or resonances, for example at period 9 for primates, vertebrates and invertebrates, or period 10-11 for yeast, bacteria and archaea as shown in [3] where the peaks were remarkably related to aspects of protein structuring and folding. Over the years, these methods and results have been extended in various ways [4], such as wavelet-type analysis [5, 6, 7] of the sequences, using features of the spectra to classify and cluster genomes with the aid of neural networks [8], prediction of coding regions [9] and periodic structures [10], etc.

In this paper, we study the Fourier spectral density of the genome of coronaviruses — a positive-sense single-stranded RNA genome with size ranging from roughly 26 to 32 kilobases, based on the dataset of [11] which covers all four genera of coronaviruses. In addition, motivated by the recent COVID-19 pandemic, we include the genomes of SARS-CoV-2, a bat coronavirus Bat-RaTG13 of close genome identity, and the MERS coronavirus.

Across the 30 different genome sequences, we find that their Fourier spectra take on the same form. There is a low frequency domain (*k* ≲ 10 in units of inverse genome length) where a sinc-squared-like oscillatory form is enveloped by a roughly 1*/k*^2^ decay curve. This is followed by stochastic white-noise type fluctuations about fixed mean values which tend to vary inversely with the genome size. We find that a random, uncorrelated sequence — with the probability of occurrence for each nucleotide being its frequency ratio in the sequence — yields similar behavior in the low-frequency domain. We develop a few models to characterize the typical spectrum, and in the process stumble upon a linear scaling law between a measure of the complexity of each genome and the power-law exponent that describes the enveloping curve of the low-frequency domain. The complexity measure that we use here is intimately related to the Shannon entropy of the sequence, and thus this relation concretely realizes a way by which information-theoretic content is carried within the genome’s spectral density.

Now, power-law decay of the form 1*/k*^*α*^ have previously been discussed in literature for other types of genomes (see for example [12, 13]). We would like to emphasize that here, we do not employ either the Fast Fourier Transform or non-overlapping averaging procedures to smoothen the data in the low-frequency domain. These are common techniques used for easing computations in past related works, but may compromise the sensitivity by which we characterize the spectral curves. We also perform the spectral density analysis at the level of the coding region (a few thousand nucleotides) for the Spike protein, an essential protein that binds to the host cell’s receptor. We find that interestingly, the general features of the spectrum persist at the protein level, but not the scaling law mentioned above.

Our paper is organized as follows. In Section 2, we present some background theory for our work, followed by Section 3 where we present the results and a few graphical plots for visualization, before concluding in Section 4. The Appendix A collects a table listing all the GenBank accession numbers [14] of the genomes, and another gathers several graphs useful for interpreting our various results.

## 2 Theoretical preliminaries

In this Section, we present some essential mathematical concepts that form the basis for our study. Our analysis of the RNA genomes can only begin after transformation of the genome sequence consisting of the four nucleotides (Adenine, Cytosine, Guanine and Uracil) into a numerical string. The spectral density of interest here is the absolute square of the discrete Fourier transform of a nucleotide indicator function *ϕ*(*i*) defined as follows

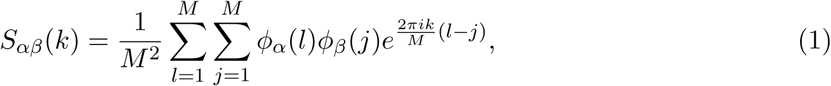

where *M* denotes the length of the genome, and *α, β* denote particular choices of nucleotides. In the continuum limit and after averaging over some distribution of genomes, this approaches the Fourier transform of the correlation function 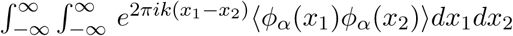.

Now, a basic premise lies in the choice of the indicator function *ϕ*(*i*). While various propositions have been explored in the literature, in this paper, following [2], we use a simple binary-valued model where for each nucleotide, *ϕ*(*i*) is equal to 1 if the nucleotide is found at position ‘*i*’ and 0 otherwise. For all our genome data, we find that (1) exhibits a clear specific oscillatory form that resembles a sinc(-squared) function in the low-frequency domain (up to *k* ∼ 10). In the following, we furnish a potential simple explanation of such low-frequency behavior. For simplicity and definiteness, we will mainly focus on the spectral density sum

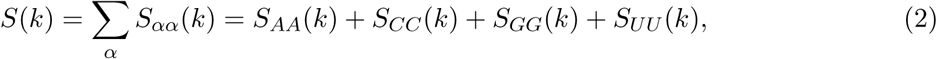

for the rest of the paper, but we have checked that the general features described above pertain to the cross-spectra *S*_*αβ*_ (with *β* ≠ *α*) as well as the individual autocorrelations *S*_*αα*_ for all the four nucleotides.

Apart from computing various quantities at the level of the entire RNA genome, we also examine the spectral density associated with the coding region for the Spike protein. For the coronaviruses, apart from the Spike protein, the genome encodes several proteins each carrying unique functions, such as the envelope, membrane, nucleocapsid, etc. In particular, the Spike protein plays an essential role in host cell receptor binding during the process of viral infection, and is thus a common target for developments of antibodies and vaccines (see for example, [11]). Now the coding region associated with this protein is only of the order of 10^3^ nucleotides, so a priori it is not clear if the spectral density can be meaningfully analyzed. We find however that the general features of the spectral density persist for the Spike protein’s coding region too.

### 2.1 A reference curve: the uncorrelated background

Consider the case of an uncorrelated numerical sequence, where the probablity of a nucleotide of type *α* occurring at some position is a constant, independent of others and the position itself. Given *N*_*α*_ such nucleotides in the sequence, we can estimate this constant to be 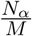, with the expectation value of the spectral density being

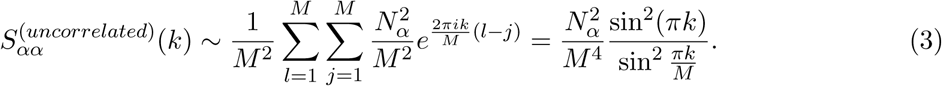

We would find that up to *k* ∼ 10, (3) models the spectral density rather well. For the local maxima of (3), they approximately occur at half-integer values of *k* and thus the upper envelope of the oscillation is manifest as a 1*/k*^2^ decay function in this domain which follows from expanding (3) about *k* = 0. The decaying behavior of the envelope curve typically stops at about *k* ∼ 30, and thereafter the spectral density appears to be characterized by stochastic fluctuations about some fixed mean.

Although (3) appears to model observed datasets well, the goodness of fit doesn’t extend beyond the *k* ∼ 10 range, nor is it clear from the data whether deviations from (3) are unimportant random fluctuations or otherwise within the low-frequency domain. To gain further insights, we present a few simple models which characterize the observed deviations from (3). The models’ parameters can potentially be used for clustering coronavirus genomes if future studies prove that these values persist for a larger sets of data, or more interestingly, they could potentially demonstrate correlation with other features of the genome that would help us recognize the presence of long-range correlations. From now on, we refer to (3) as the ‘uncorrelated background’.

### 2.2 Three simple models

In the following, we present three models for the observed spectral density that characterize deviations from the uncorrelated background. The first two concerns the description of the low-frequency domain (*k* ≲ 10) whereas the third involves a more global description.

#### (A) Power-law decay of the enveloping curve

Motivated by previous works on this subject, we consider fitting a power-law decay via least-square regression to the enveloping curve (for *k* ∈ [1, 10]) of the form

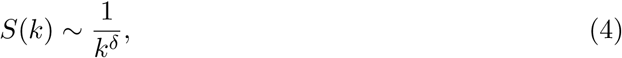

for some power exponent *δ*. The power-law description is convenient and has proven to be a popularly studied model for spectral density of genomes in general (see for example [13]). It is crucial to bear in mind that it is a coarse-grained description which doesn’t extend to the origin, and valid only for the low-frequency domain. We would later find that this is the parameter that remarkably scales linearly with a measure of the genome complexity. For all our datasets, *ϵ* = *δ* − 2 ∼ 10^−2^. It is not a priori clear how large *ϵ* has to be in order for the deviation to be significant, and more sequences corresponding to each type of coronavirus should be studied in order to determine the range of *ϵ* and its statistical distribution. Although we leave this for future work, we found evidence that the variation in the *ϵ* correlates with a measure of the complexity of the genome (which at the limit of infinite genome size approaches the Shannon entropy) in a way that is distinctly different from a completely random sequence.

It is useful to compute the expected *δ* for the hypothetical uncorrelated background (3) which is parametrized by the genome size *M* and the sum of squares of nucleotide number 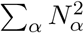. For the general spectral density *S*(*k*), from least-square regression of the log-log relation, we obtain *ϵ* ≡ 2 − *δ* to be

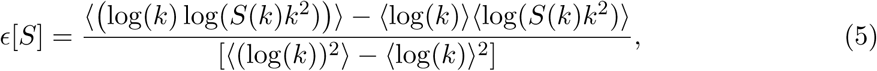

where 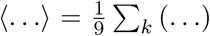 denotes averaging over the nine local maxima points in the domain *k* ∈ [1, 10]. For the uncorrelated case of (3), we find that the factor 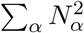 cancels away in (5) and numerically, *δ* ≈ 1.956 for all the datasets at the level of the genome and that of the protein coding region. This defines a background value for the detection of a deviation away from the completely random sequence.

#### (B) Linearized correlation function

In contrast to an empirical power-law fitting of only the enveloping curve, one could adopt a bottom-up approach by postulating certain forms of the correlation function, and then performing the discrete Fourier transform. Consider the case where the correlation function is a linear function of the nucleotide separation, we can write

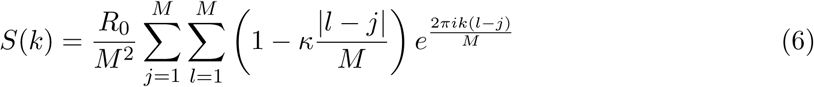

for some constant *κ*, and 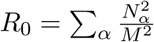. A straightforward calculation yields

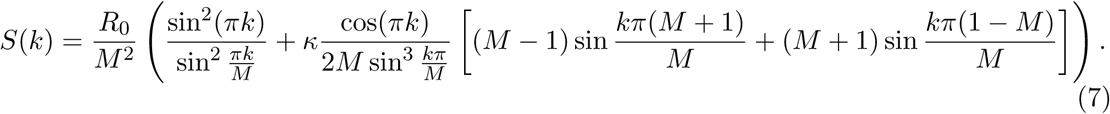

This function is invariant under the reflection *k* ↔ *M* − *k*, which is an exact discrete symmetry for the spectral density *S*(*k*) (or the individual *S*_*αα*_(*k*)) more generally. The parameter *κ* admits the physical interpretation of the presence of long-range correlation/anti-correlation depending on whether it’s positive/negative, and we would find that apart from one exception, all our datasets can be matched to a positive *κ* of the order 10^−2^. We find that if the curve-fitting is performed taking into account only the first ten local maxima as in the case for *δ*-parameter, the local minima points at integral *k*-values are not well captured by the fitted curve, so we also include them in the curve-fitting.

Beyond the specific linear form of the correlation function postulated in (6), it is also representative of a large class of correlation functions of the form

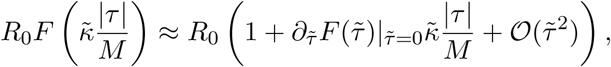

where *τ* ≡ *l* − *j*, 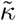 is a small constant and 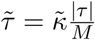. This first-order truncation is identical to (6) with 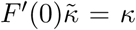. Thus, (6) could approximate correlation functions of the general form 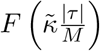 where 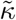 is a small dimensionless parameter, and

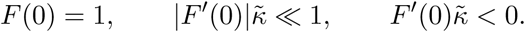

For example, if the correlation function turns out to be an exponentially decaying function of the form 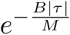 with *B* ≪ 1, then to a good approximation we can identify *κ* ∼ *B*.

#### (c) A Lorentzian function

The power-law decay in (A) parametrizes the decay of the envelope whereas the model in could account for non-vanishing local minima in the low-frequency domain. Beyond this region, we seek an interpolating curve that extends throughout the spectrum including the origin. For this purpose, we consider fitting a Lorentzian function of the following form to the spectrum

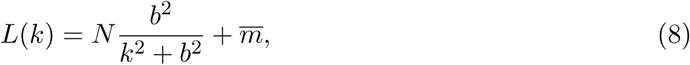

where 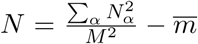 and 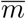 is the mean value near the spectrum’s midpoint, about which stochastic fluctuations are observed.^2^ This is a simple coarse-grained model which averages over the oscillations in the low-frequency domain and describes the overall decay of the spectrum via a smooth curve. Like the *κ* parameter in the model (6), the curve-fitting is performed with the set of extremal points in the low-frequency domain, with the initial and final conditions taken into account by first fixing *N*, 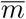 with their observed values for each genome sequence. As a useful reference, we also fit the Lorentzian function to the uncorrelated background (3) and finding *b*^2^ ≈ 0.0765 with 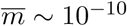 at the genome level, and 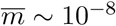 at the protein coding region level.

### 2.3 A measure of complexity and Shannon entropy

Scaling laws manifest in the Fourier spectral density have often motivated the study of features of the genome that reflect various properties of it being a complex system, such as the fractal dimension (of a suitably defined matrix representation of the correlation function), etc. A measure of the complexity of the genome considered in the past literature (see for example [15, 16, 17]) is defined as follows.

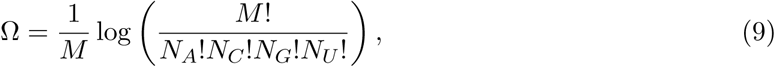

where *N*_*α*_ is the number of the *α*-nucleotide. The logarithmic argument counts the number of distinguishable permutations given a fixed number of each nucleotide. At large *M*, this admits a natural interpretation of the Shannon entropy of the genome sequence. To see this, we can invoke Stirling’s formula to express the large-*M* limit of Ω as

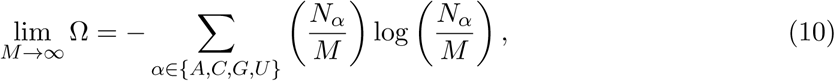

which is a function of only the fractional distribution of nucleotides. In this form (10), the measure of complexity Ω is clearly the Shannon entropy which measures the information entropy associated with a genome sequence where the probability of nucleotide-*α* occurring in any position is 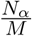. We would find later that interestingly, the model parameter *δ* (but not *κ*) scales linearly with Ω across the dataset of 30 types of coronavirus genomes. Also, when restricted to the Spike protein’s level, the measure of complexity appears to scale linearly with the overall measure at the genome level. But the model parameter *δ* that is computed at the level of the Spike protein does not correlate with Ω at either the genome/protein level, and neither does *κ*.

## 3 Results and graphs

Our genome dataset^3^ consisting of 30 types of coronaviruses spread across four genera mainly follows from reference [11] plus a few other additions: SARS-CoV-2, MERS-CoV and Bat-RaTG13. Bat-RaTG13 is a bat coronavirus that was most recently found to have 96% genome identity with SARS-CoV-2 and featured in papers discussing a possible bat origin of the latter [18]. We included it here to see how the model parameters for this genome compare to that of SARS-CoV-2 relative to the other coronaviruses. In the following, we outline the essential results, using the example of the SARS-CoV-2 reference genome for various graphical illustrations.

We find that the Fourier spectral density is characterized by the following features:

a. In a small low-frequency regime (*k* ≲ 10), the uncorrelated background (3) is a good approximation (see Fig. 1) for all genome sequences we examined. After curve-fitting to the datasets, we find the following range of values for the model parameters:^4^

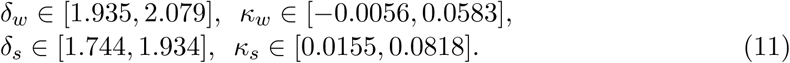

From visual inspection of the relevant graphical plots, we find no obvious correlation among these model parameters, nor between them and the genome/Spike protein sizes. But we find that *δ*_*w*_ and Ω_*w*_ appear to be related. Linear regression yields the following best-fit line (see Fig. 2)

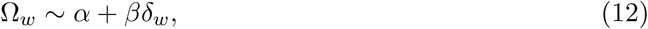

**Figure 1:**
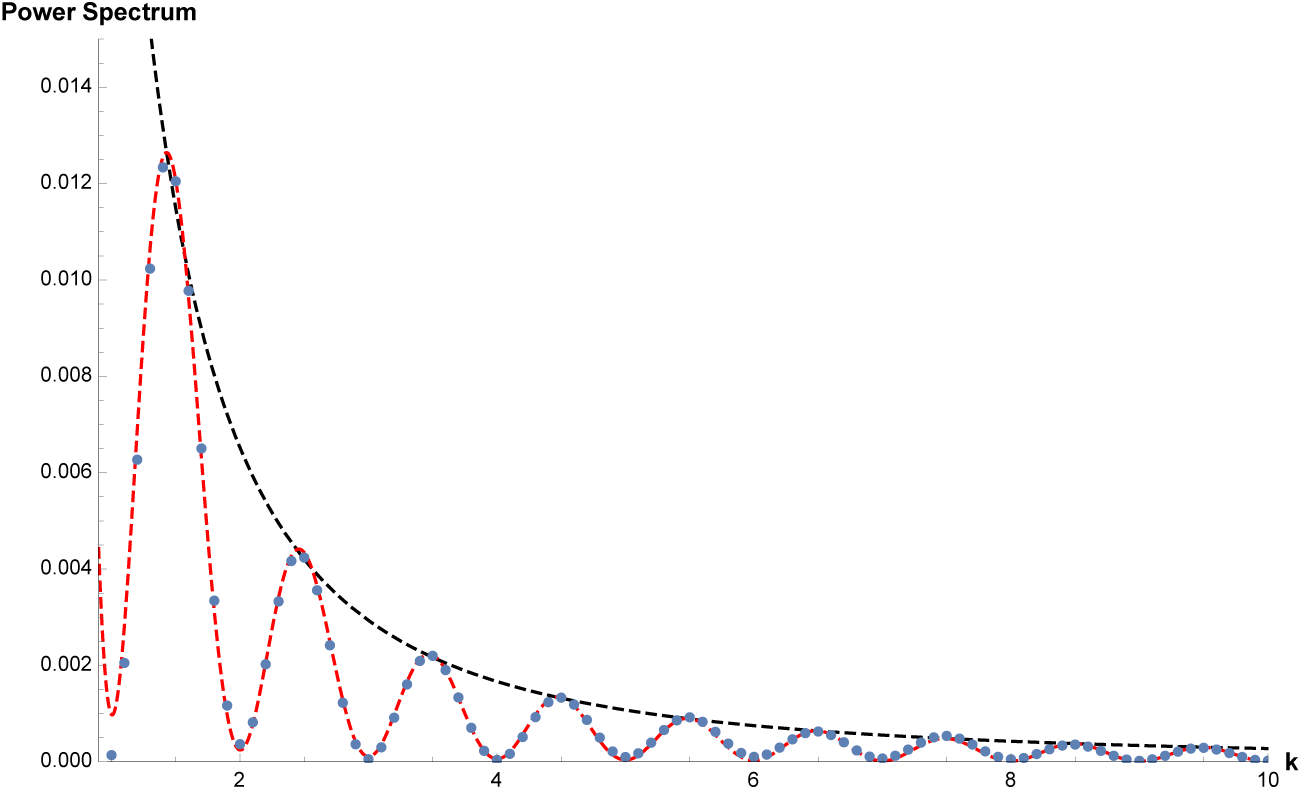
Plot of the power spectral density (SARS-CoV-2 genome) in the range where we perform the curve-fitting. The black dashed line is the curve ∼ 1*/k*^*δ*^, *δ* = 1.968 obtained from the set of local maxima, while the red dashed line is equation (7) obtained by fitting to all maxima and minima, with the best-fit value *κ* = 0.0362.

**Figure 2:**
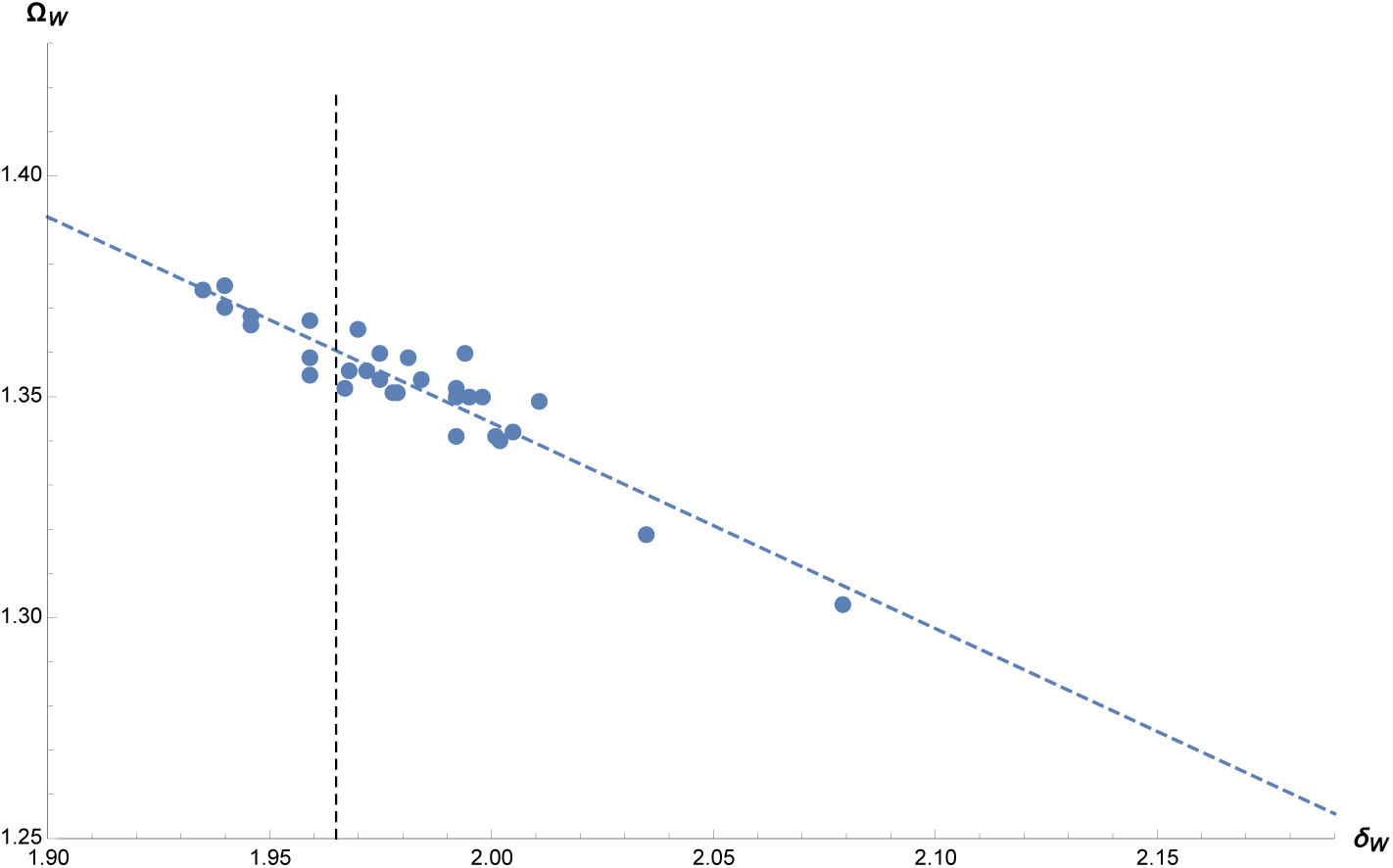
Plot showing linear regression fit for (*δ*_*w*_, Ω_*w*_) parameters. In the absence of any correlation, we would instead observe a vertical line at *δ*_*w*_ = 1.956 — the value that corresponds to (3). with the line parameters being (with the 95% confidence intervals in brackets)

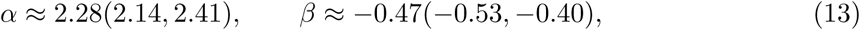

Since we checked that for all the 30 coronaviruses, the assumption of a completely uncorrelated background yields *δ* ≈ 1.956, this leads to a convenient definition of a reference complexity value

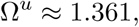

which lies at the intersection between the uncorrelated vertical line and the observed one with finite slope. The difference between the observed complexity measure and Ω^*u*^ in turn enacts a measure of the deviation from complete randomness of the sequence. There is also a similar relation between *δ*_*w*_ and Ω_*s*_, consistent with the following linear relation that we found:

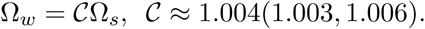

It would be interesting to study this for other coding and non-coding regions as it is suggestive of some level of self-similarity for this complexity measure.
b. After *k* ∼ 10, the genome displays much more scatter about the uncorrelated background, and the models of deviation are no longer effective descriptions (see Fig. 3). Stochastic fluctuations about a fixed mean appear to set in and there are no isolated peaks apart from two prominent ones at 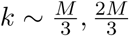 which have been seen and interpreted in past literature [2, 19] to correspond to the universal triplet codon usage. We applied an (overlapping) moving average (of window size ∼ 100 nucleotides) to smooth out the data, and checked that there is no apparent regime where some non-trivial scaling law holds (see Fig. 4 and 5). At the level of both the genome and protein coding region, the fixed mean parameter 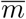 appears to correlate with the genome size. It appears to generally decrease with the size of the sequence,at both levels of the genome and the Spike protein (see Figures 6a and 6b) in Appendix B). At the genome level, it is of the order ~ 10^−5^ which is about ∼ 10^5^ larger than the value expected for the uncorrelated background, whereas at the spike protein level, 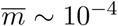 which is 10^4^ times larger than the uncorrelated background. The Lorentzian function that is fitted to the data with initial and final conditions fixed by *R*_0_ and 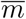 is parametrized by the half-width parameter *b*. We find that this parameter generally increases with *κ* at both genome and Spike protein levels (see Figures 7a and 7b in Appendix B).

**Figure 3:**
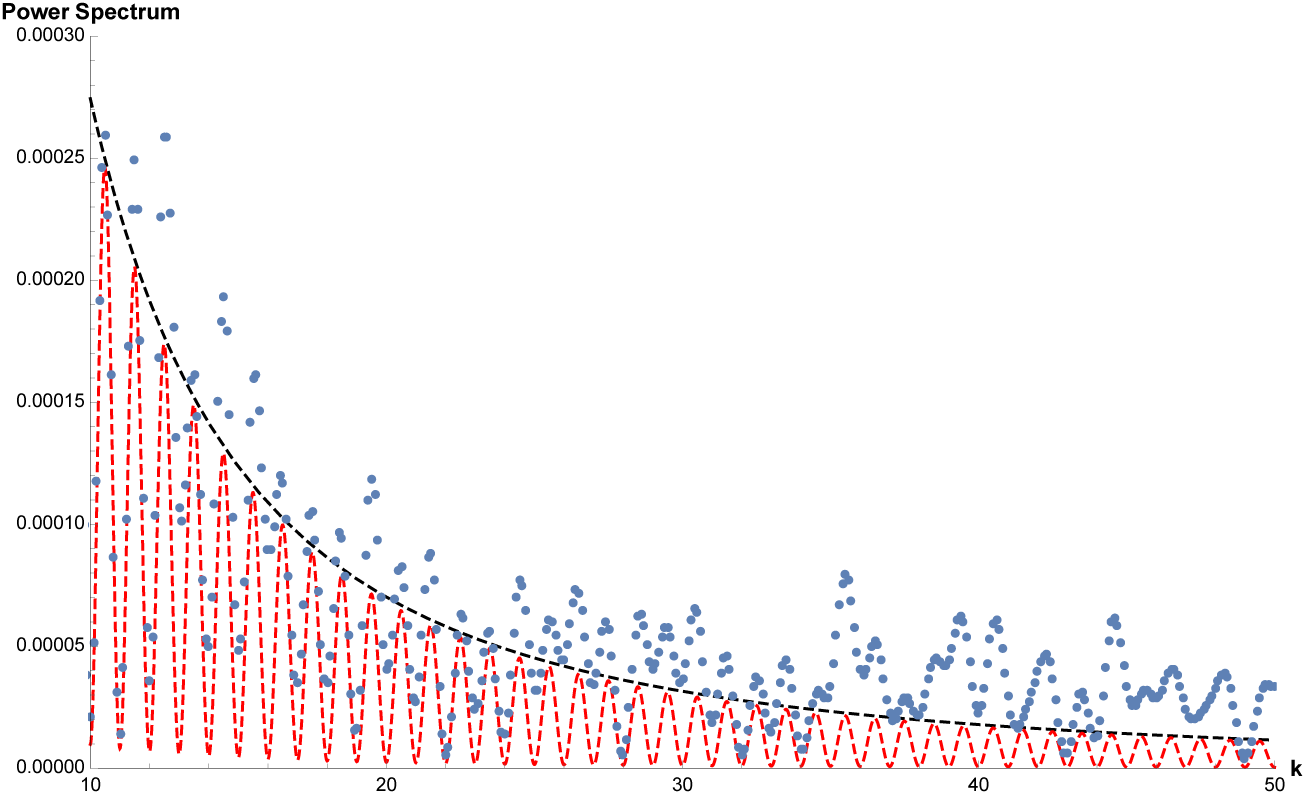
Plot of the spectral density (SARS-CoV-2 genome) showing how after about *k* = 10, the data points appear to be noisy and such stochastic fluctuations appear to persist throughout apart from a couple of isolated peaks. Neither the envelope curve of 1*/k*^*δ*^ nor equation (7) continue to be effective descriptions.

**Figure 4:**
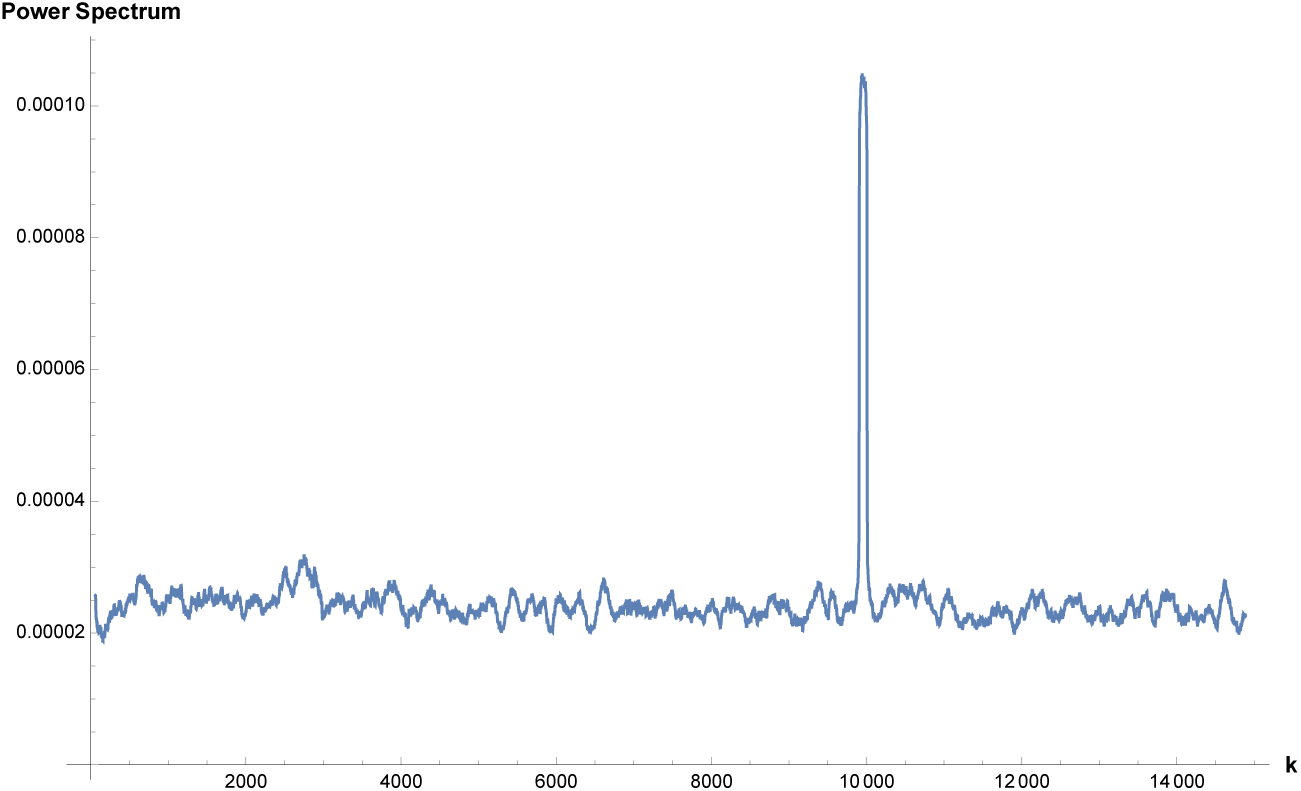
The smoothened data presents a stochastic fluctuation about a fixed mean ∼ 2×10^−5^ and there is only an isolated peak at *M/*3 due to the triplet codon-usage. Only half of the spectrum is shown here since the other half is a reflection of it due to the discrete symmetry *S*(*k*) ↔ *S*(*M* − *k*) the spectrum as mentioned in Section 2.

**Figure 5:**
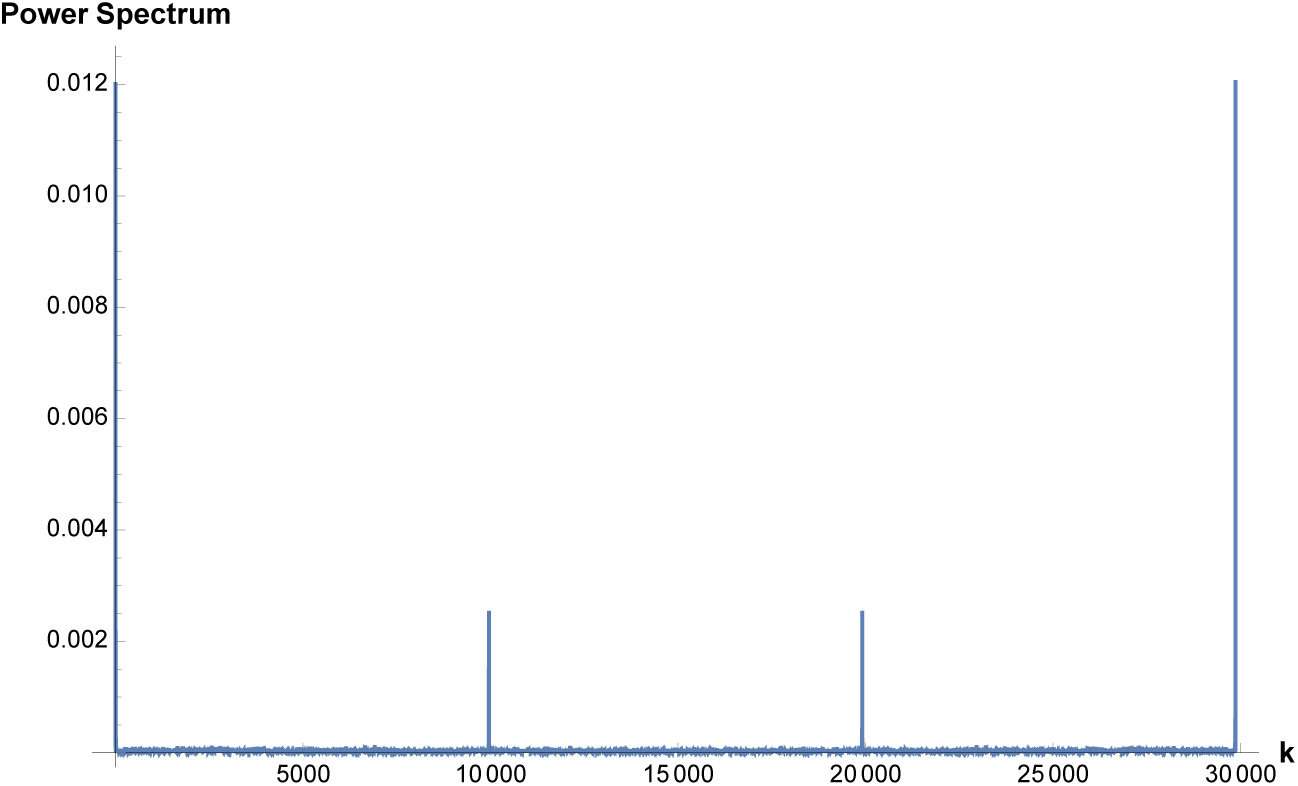
Plot of the Fourier spectral density (SARS-CoV-2 genome) which is mostly featureless with noise apart from prominent peaks at 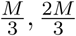 which correspond to triplet-codon usage.

**Figure 6:**
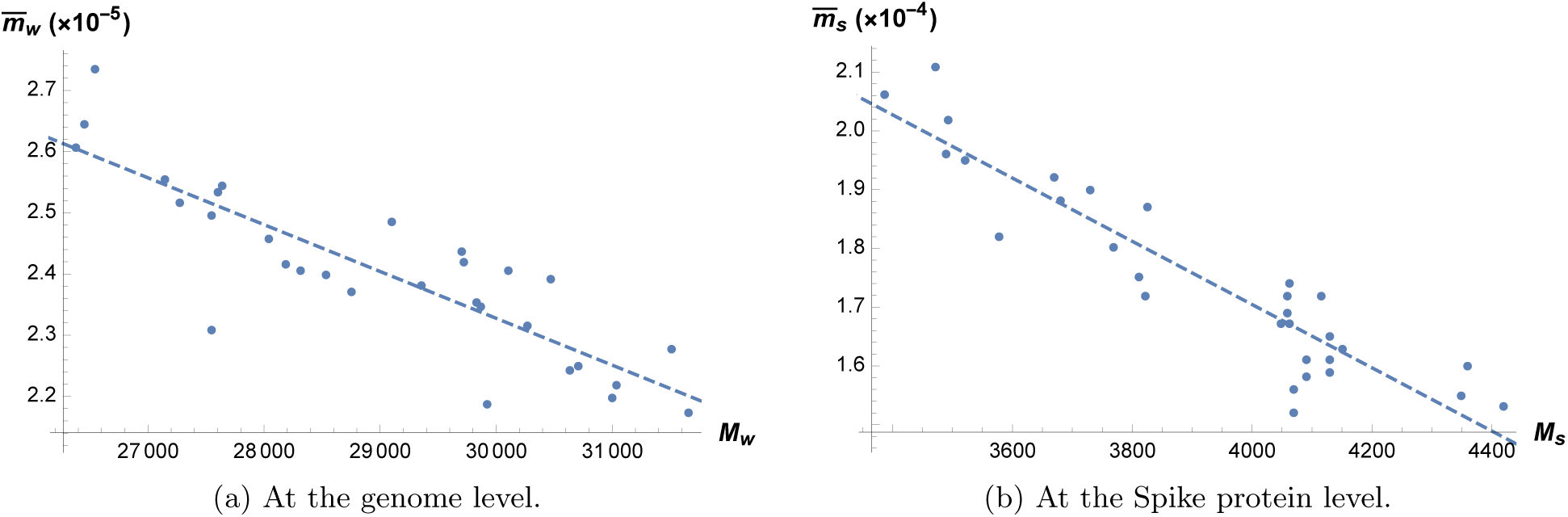
Plots showing how 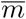 generally decreases with sequence size at both genome and Spike protein levels. The dashed line is obtained from least-square regression. For the genome, we have 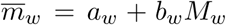, with *a*_*w*_*/*10^−5^ ≈ 4.62(4.11, 5.14), *b*_*w*_*/*10^−10^ ≈ −7.65(−9.42, −5.89), whereas for the Spike protein, we have 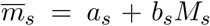, with *a*_*s*_*/*10^−4^ ≈ 3.86(3.52, 4.19), *b*_*s*_*/*10^−8^ ≈ −5.38(−6.24, −4.52).

**Figure 7:**
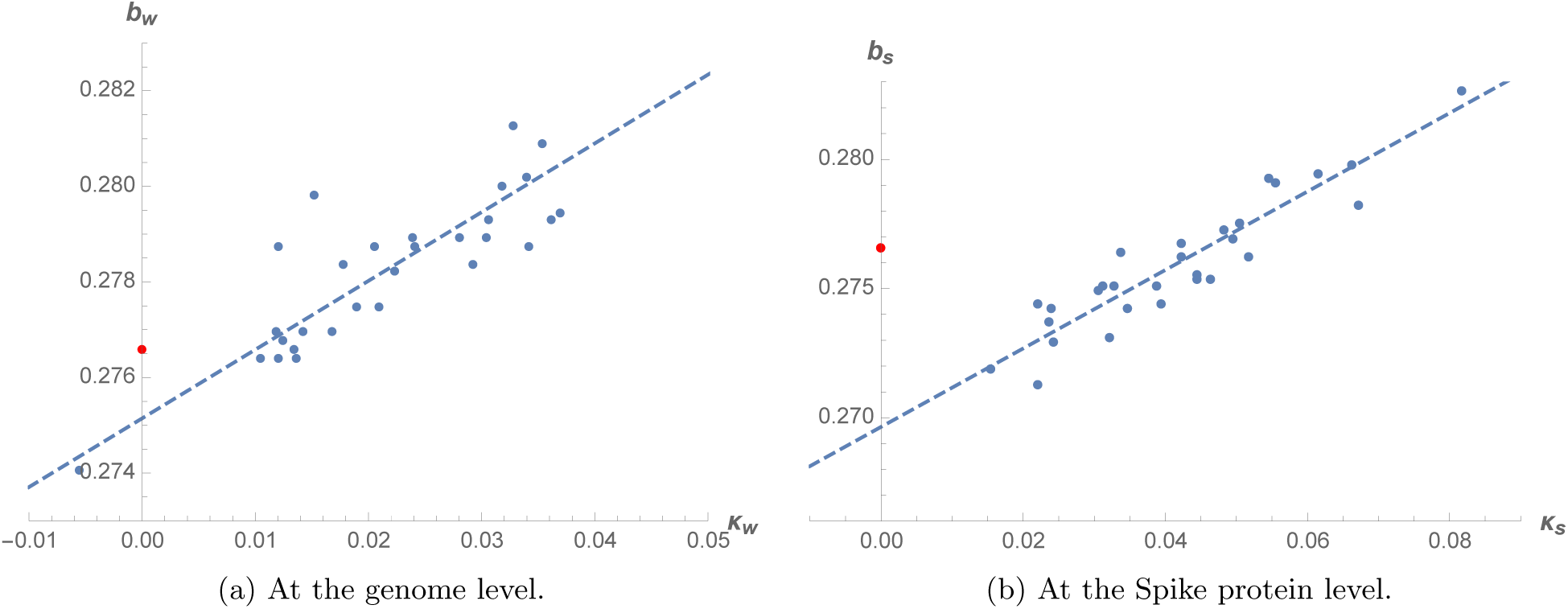
Plots showing how the half-width parameter *b* generally increases with *κ*. At the genome level, the best-fit line is *b*_*w*_ = *g*_*w*_ + *h*_*w*_*κ*_*w*_ with *g*_*w*_ ≈ 0.275(0.274, 0.276), *h*_*w*_ ≈ 0.144(0.116, 0.172). At the Spike protein level, the best-fit line is *b*_*s*_ = *g*_*s*_ + *h*_*s*_*κ*_*s*_, with *g*_*s*_ ≈ 0.270(0.269, 0.271), *h*_*s*_ ≈ 0.152(0.128, 0.176). The red point pertains to the uncorrelated background.
c. Finally, although for simplicity, we have kept to analyzing the spectral density corresponding to the sum of all the nucleotides, the general qualitative features described in (a) and (b) above apply to the spectral density for each individual nucleotide as well as the cross-spectra.

## 4 Discussion

We have presented a study of the Fourier spectral density of the coronavirus genome at the level of the entire genome as well as the coding region for the Spike protein. The power spectrum profile can be well-described by considering aspects of deviation from the hypothetical case of a random, uncorrelated sequence (eqn. (3)). We summarize the essential general features below:

i. There is a low-frequency domain (*k* ≲ 10) which exhibits a clear oscillatory form that is close to (3). In this domain, we find that the enveloping curve connecting the local maxima is well-described by a power decay law of the form 1*/k*^*δ*^. We noted that the power exponent *δ* shows a correlation with a measure of complexity of the sequence (eqn. (9)) which in the limit of large genome size is the sequence’s Shannon entropy. The deviation from the uncorrelated background can be described by a linear relation between *δ* and Ω. This behavior does not however persist at the level of the Spike protein’s coding region.
ii. Beyond the low-frequency domain, the spectrum displays stochastic fluctuations about certain fixed values 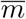, and we find no other resonances apart from the peaks at 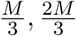 which are associated with the universal triplet codon usage. Relative to the uncorrelated case, 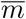 is about 10^5^ higher at the genome level and about 10^4^ higher at the Spike protein level. It also generally decreases with the size of the genome or the protein coding region.
iii. Upon fitting the Lorentzian function to the spectrum with initial and final conditions determined by *R*_0_ and 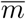 respectively, we find that its half-width parameter is correlated with *κ* — the dimensionless constant that defines the linearized correlation function in the low-frequency domain, and generally increases with it. This is observed at both the genome and Spike protein’s levels.

Let us conclude by briefly pointing out several future directions and applications. Now, it has been noted in literature for some time that DNA viruses and unicellular organisms tend to have mutation rates which vary inversely with the genome size (‘Drake’s rule’ [20, 21, 22]). This correlation has been studied for RNA viruses recently (see for example [23]) although we are unaware of any evidence for the case of coronaviruses^5^ which is the only RNA virus family which has a 3’-exonuclease proofreading mechanism that enhances replication fidelity. The parameter 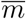 that we have introduced here appears to vary inversely with genome size, and thus it may be worthwhile to explore its role in models that attempts to explain viral mutation rates. In [25], a negative association between molecular evolution rate and genome size was established for RNA viruses. It would be interesting to compute the parameter 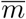 for the viral sequences studied in [25, 26]. Another potential application of our work which has immediate relevance is to study the distribution of 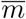 for SARS-CoV-2 genomes specifically to explore if they could describe current evolution of the virus (see for example [27]).

The Lorentzian function that we fit broadly to the spectrum as a whole is a coarse-grained description that does not model the transition from the low-frequency spectrum to the other part of the spectrum that appears to be dominated by stochastic fluctuations. It would be interesting to develop theoretical models that could possibly account for such a transition and in the process, construct a clearer understanding for the parameter 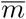 or why the information-theoretic measure (9) is relevant for the low-frequency domain.

A complementary approach towards understanding correlation effects is to study directly the correlation function itself (see for example [28]), although this is more computationally intensive. It would be interesting to study what forms of correlation functions could lead to the enveloping curve being of the form 1*/k*^*δ*^. A few related models were proposed in [29, 30], and it may be worthwhile to revisit them in light of the newfound relation with the measure of complexity.

Finally, it would be interesting to perform a more extensive study of the models here with a larger set of viral genomes so that we have a fuller understanding of their statistical distribution and whether they can be useful in clustering and classifying purposes.^6^ Motivated by the COVID-19 pandemic, notwithstanding our limited dataset, in Table 1 below, we show the viral genome that is the closest neighbor to SARS-CoV-2 for each of the four model parameters at both levels of the genome and Spike protein coding region. From Table 1, we see that Bat-RTG13 features most frequently and that apart from TGEV and HKU1 which infect pigs and humans respectively, the others are bat coronaviruses. Collectively, they appear to be broadly compatible with the plausibility of the bat origin of SARS-CoV-2, while to our knowledge, the association of SARS-CoV-2 with TGEV and HKU1 has never been made in literature.

**Table 1:**
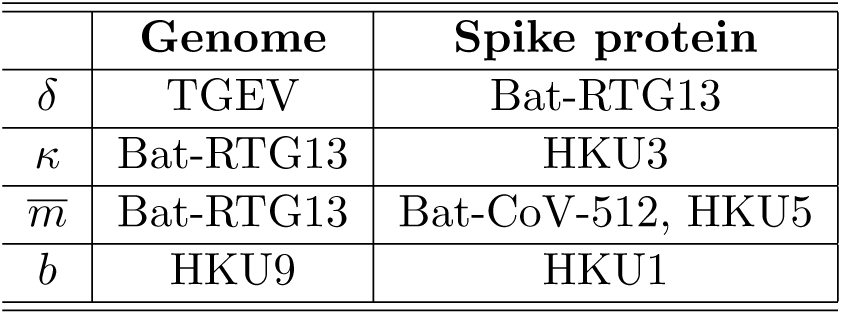
We display the coronavirus that has a genome closest to SARS-CoV-2 in terms of each of the model parameters.

|  | **Genome** | **Spike protein** |
| --- | --- | --- |
| $\delta$ | TGEV | Bat-RTG13 |
| $\kappa$ | Bat-RTG13 | HKU3 |
| $\overline{m}$ | Bat-RTG13 | Bat-CoV-512, HKU5 |
| $b$ | HKU9 | HKU1 |

## Acknowledgments

I thank Neal Snyderman and Rajesh Parwani for stimulating discussions.

## A GenBank accession numbers

This appendix collects the GenBank accession ID and names of the 30 coronaviruses used in this work, which largely follows [11], our additions being SARS-CoV-2, MERS-CoV and Bat-RaTG13. These genomes can be freely downloaded from https://www.ncbi.nlm.nih.gov. For each genome, we exclude the poly(A) tail for our analysis.

**Table 2:** The GenBank accession ID for the thirty coronavirus RNA genomes covered in this work.

| Virus | GenBank accession number |
| --- | --- |
| *Alphacoronavirus* |  |
| Transmissible gastroenteritis virus (TGEV) | DQ811785 |
| Porcine respiratory coronavirus (PCRV) | DQ811787 |
| Feline coronavirus (FCoV) | NC_002306 |
| Human coronavirus 229E (HCoV-229E) | NC_002645 |
| Human coronavirus NL63 (HCoV-NL63) | NC_005831 |
| Porcine epidemic diarrhea virus (PEDV) | NC_003436 |
| Scotophilus bat coronavirus 512 | NC_009657 |
| Rhinolophus bat coronavirus HKU2 | NC_009988 |
| Miniopterus bat coronavirus HKU8 | NC_010438 |
| Miniopterus bat coronavirus 1A | NC_010437 |
| *Betacoronavirus* |  |
| Human coronavirus OC43 | NC_006213 |
| Bovine coronavirus (BCoV) | NC_003045 |
| Porcine hemagglutinating encephalomyelitis virus (PHEV) | KY419103 |
| Equine coronavirus (ECoV) | EF446615 |
| Human coronavirus HKU1 | NC_006577 |
| Mouse hepatitis virus (MHV) | AC_000192 |
| SARS coronavirus (SARS-CoV) | NC_004718 |
| SARS coronavirus-2 (SARS-CoV-2) | NC_045512 |
| Bat SARS coronavirus HKU3 | GQ153539 |
| Bat coronavirus RaTG13 | MN996532 |
| MERS-coronavirus | NC_019843 |
| Tylonycteris bat coronavirus HKU4 | NC_009019 |
| Pipistrellus bat coronavirus HKU5 | NC_009020 |
| Rousettus bat coronavirus HKU9 | NC_009021 |
| *Gammacoronavirus* |  |
| Infectious bronchitis virus (IBV) | NC_001451 |
| Beluga whale coronavirus (SW1) | NC_010646 |
| Turkey coronavirus (TCoV) | NC_010800 |
| *Deltacoronavirus* |  |
| Munia coronavirus HKU13 | NC_011550 |
| Thrush coronavirus HKU12 | NC_011549 |
| Bulbul coronavirus HKU11 | FJ376620 |

## B Some Graphs

In this Section, we collect several graphs useful for visualizing two particular trends observed: (i) the parameter 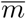 tends to vary inversely with size of genome/Spike protein coding region, (ii) the linearized correlation function parameter *κ* and the half-width parameter *b* appears to be correlated.

## Footnotes

1 See for example [1] for an inspiring read.

2 We only consider half of the spectrum, since the other half naturally arises from the discrete symmetry *S*(*k*) = *S*(*M* − *k*).

3 We list their names and GenBank accession IDs of the genomes in the Appendix for reference.

4 Whenever appropriate, we use the subscripts to label the level (*w* = whole genome, *s* = Spike protein) at which various quantities are computed.

5 Some recent results concerning mutation rates of coronaviruses were published in [24].

6 See [31] for a recent attempt in this direction for coronaviruses.

